# Harzianic Acid has multi-target antimicrobial activity against Gram-Positive Bacteria

**DOI:** 10.1101/2021.12.03.471066

**Authors:** Xudong Ouyang, Jelmer Hoeksma, Wouter A.G. Beenker, Samantha van der Beek, Jeroen den Hertog

## Abstract

The thermophilic fungus *Oidiodendron flavum* is a saprobe that is commonly isolated from soil. Here, we identified a Gram-positive bacteria-selective antimicrobial secondary metabolite from this fungal species, harzianic acid (HA). Using *Bacillus subtilis* strain 168 combined with several assays, we found that HA targeted the cell membrane, though only at high concentrations. To further study the antimicrobial activity of HA, we isolated an HA-resistant strain, *Bacillus subtilis* strain M9015, and discovered that the mutant has more translucent colonies, has cross resistance to rifampin, and harbors five mutations in the coding region of four distinct genes. Further analysis of these genes indicated that the mutation in *atpE* might be responsible for the translucency of the colony, and mutation in *yusO* for resistance to both HA and rifampin. We conclude that HA is a multi-target antimicrobial agent against Gram-positive bacteria.

## Introduction

Fungi interact with their surroundings by production and secretion of biologically active compounds, secondary metabolites (SMs), which are not essential for fungal growth (1). These compounds are chemically distinct small molecules (in most cases < 3 kDa) often with biological activities that are produced at specific stages of growth to perform important functions, including survival from harsh environments, communication with invaders or alteration of fungal development (2). SMs are synthesized along different pathways than primary metabolites (3) and they are excellent sources of potential therapeutic drugs (4).

The genus of *Oidiodendron* was established under the *Myxotrichaceae* family by Robak in 1932 (5). Species of *Oidiodendron* are known as saprobes and are commonly isolated from a wide range of habitats, including soil, decaying plant materials, marine sediments and decomposing human hair (6-9). They primarily occur through the temperate regions, with a few exceptions from tropical and subtropical locales. The widespread distribution of this genus is in connection with their excellent adaptive capacity, by which they establish various interactions with other organisms (10-12). Whereas the genus of *Oidiodendron* has drawn much attention with respect to studies of morphology and taxology (10, 13), there are not many investigations into their ability to produce SMs.

Previously, in order to search for novel bioactive compounds, we have established a fungal SMs library containing more than 10,000 fungal species (14), including several *Oidiodendron* strains. One of these strains, *Oidiodendron flavum*, was found to produce antimicrobial activity, which was identified as harzianic acid (HA) (15). HA was first isolated as a novel antimicrobial agent from a fungal strain *Trichoderma harzianum* in 1994 (16). Surprisingly, subsequent research focused more on the activity of HA as a plant promotor rather than an antimicrobial (3, 17). Recently, more interest has been drawn back into its antimicrobial activities. Xie et al. has described HA as an inhibitor of acetohydroxyacid synthase (AHAS) in fungi (18). However in bacteria, although its activity against pathogenic bacteria has been reported (19), not much data are available on its targets yet. Since HA strongly inhibits bacterial growth, it is suggested to be a promising candidate antibacterial agent. Thus, further research into identification of its mechanism of action (MoA) is required. Here, we determined the antimicrobial spectrum of HA on 14 bacterial strains, and investigated HA’s MoA with a combination of assays. In addition, we have developed an HA resistant *Bacillus subtilis* strain by continuous exposure to HA. This strain produced translucent colonies and was four times more resistant to HA. Surprisingly, compared to wild type, this mutant strain was resistant to up to eight times higher concentrations of another antimicrobial agent, rifampin, as well. Genetic analysis of this mutant strain indicated that a recently identified multidrug resistance operon was involved in the enhanced resistance to HA.

## Results

### Identification of HA

In a screen for antimicrobial activity, SMs from *O. flavum* were identified to have potent activity. In order to identify the active compound from *O. flavum*, the agar culture of this fungus was extracted using ethyl acetate and fractionated by HPLC. The pure active fraction was then tested for its UV-Vis spectrum, which showed maximum absorbance at 244 nm and 363 nm. Next, high resolution mass spectrometry (HRMS) of the active compound revealed a mass of the sodium adduct of 388.1750, which suggested several options for a molecular formula. Finally, the remainder of the fraction was dried and used for elemental composition analyses, showing mass percentages of 61,4% carbon, 21,7% oxygen, 7,0% hydrogen and 3,8% nitrogen. Based on these analyses, we determined a molecular formula of C_19_H_27_NO_6_. Through NMR spectroscopy we obtained ^1^H, ^13^C, Heteronuclear Single-Quantum Correlation Spectroscopy (HSQC), Heteronuclear Multiple-Bond Correlation spectroscopy (HMBC) and homonuclear correlation (COSY) spectra (Table S1, Figure S1A-E), which were consistent with data previously reported for HA (3, 16) (Figure S1F). Thus, we concluded that HA is the antimicrobial compound from *O. flavum*.

**Table 1.** MICs of HA and rifampin on B. subtilis and mutants. MICs were determined by serial dilution of HA or Rifampin using the indicated strain. The result of biological triplicates is depicted here.

| Strain | Source | MIC to HA ($\mu\text{g ml}^{-1}$) | MIC to rifampin ($\mu\text{g ml}^{-1}$) |
| --- | --- | --- | --- |
| *Bacillus subtilis* strain 168 | Home stock | 50 | 0.13 |
| *Bacillus subtilis* strain M9015 | This work | 200 | 0.8 |
| *Bacillus subtilis* strain $\Delta\text{atpE}$ | BKK36860* | 25 | 0.13 |
| *Bacillus subtilis* strain $\Delta\text{ymaB}$ | BKK17400* | 50 | 0.13 |
| *Bacillus subtilis* strain $\Delta\text{yusO}$ | BKK32870* | 200 | 0.66 |
| *Bacillus subtilis* strain $\Delta\text{flgL}$ | BKE35400* | 50 | 0.13 |

\* Strains from Bacillus Genetic Stock Center.

Harzianic acid: C_19_H_27_NO_6_, dark yellow powder. HRMS: found 388.1750 (M+Na), calculated 388.1736 for C_19_H_27_NO_6_Na. Elemental composition analyses: C 61,4%; O 21,8%; H 7,0%; N 3,8%. NMR (600 MHz, CDCl3): See Table S1. UV-Vis λ_max_: 244 nm, 363 nm.

### HA is a selective antimicrobial agent against Gram-positive bacteria and induces cell lysis of Bacillus subtilis cells

HA was tested against a panel of 14 pathogenic bacteria, including seven Gram-positive and seven Gram-negative strains (Table S2). The growth of all Gram-positive bacteria was affected with minimum inhibitory concentrations (MICs) ranging from 25 to 200 μg ml^-1^. The panel included antibiotic-resistant pathogenic bacteria, which were sensitive to HA, e.g. methicillin resistant *Staphylococcus aureus* (MRSA) at 200 μg ml^-1^ and vancomycin resistant *Enterococcus faecium* (VRE) at 100 μg ml^-1^. No inhibition was observed in response to HA of any of the Gram-negative bacteria tested up to 400 μg ml^-1^ This indicates that HA is a selective antimicrobial agent against Gram-positive bacteria.

To further explore the antimicrobial properties of HA against Gram-positive bacteria, a model Gram-positive organism, *B. subtilis* strain 168, was used for the following assays. First, to obtain an accurate MIC, the growth curves of *B. subtilis* were measured in the presence of a range of HA concentrations (Figure 1A). Although cells were also affected at concentrations of 30 and 40 μg ml^-1^, total inhibition of cell growth after 18 h (overnight) was only seen at a concentration of 50 μg ml^-1^ and higher, suggesting that the correct MIC for *B. subtilis* was 50 μg ml^-1^. In the following assays, we used 250 μg ml^-1^ (5 × MIC) HA to ensure the effects on cells, unless specified differently. Interestingly, a decrease in OD_600_ was observed 30 min after addition of the compound with all concentrations of HA, even with the lowest concentration we tested, 30 μg ml^-1^. A decrease in OD_600_ was not observed with vehicle control (DMSO), indicating that the HA-induced decrease was not an artefact. These results suggest that HA might induce bacterial cell lysis.

**Figure 1.**
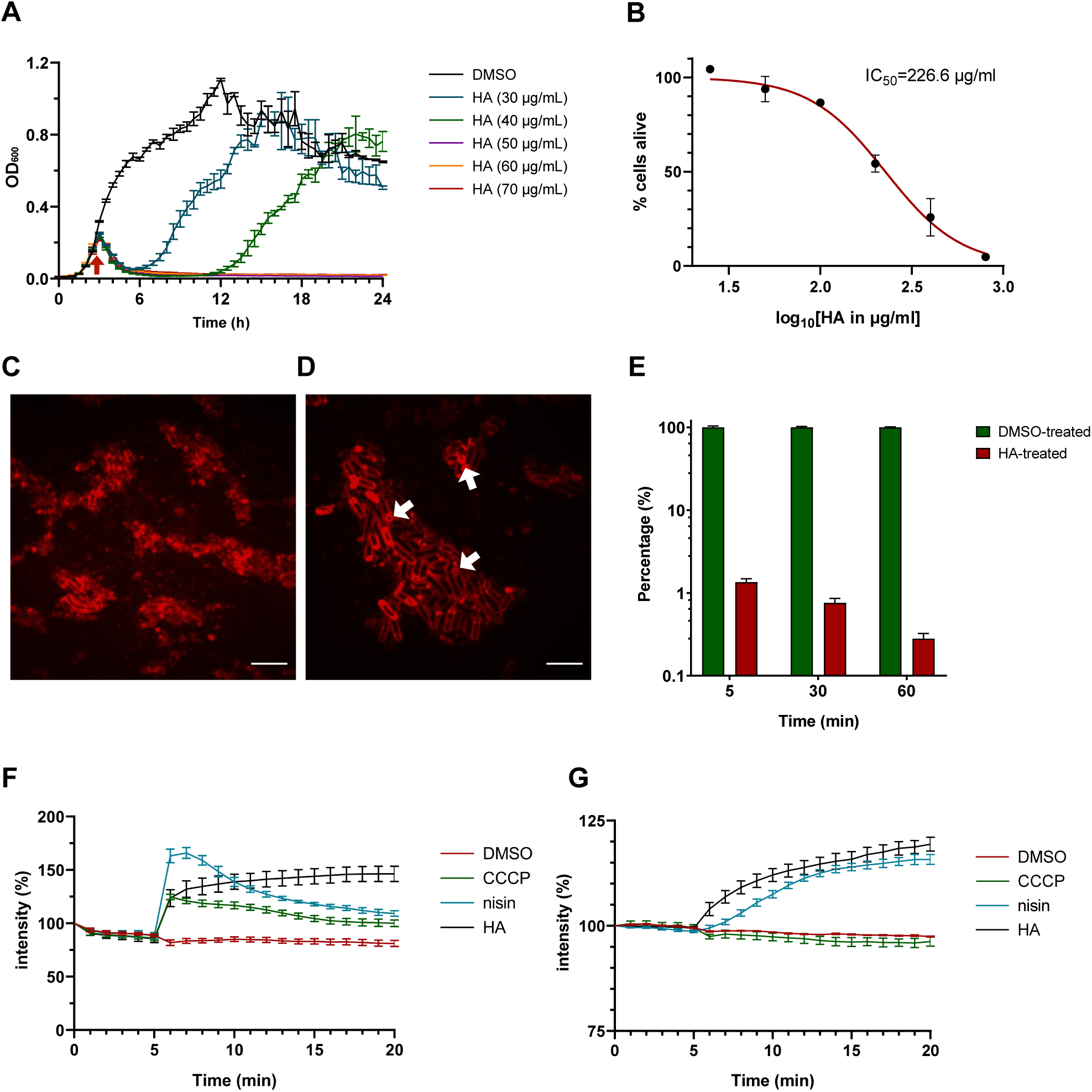
Antimicrobial property of HA. (A, B) HA inhibits eukaryotic cell growth and bacterial growth. (A) Cytotoxicity assay. HepG2 cells were incubated with HA in different concentrations for 20 h before addition of resazurin. The ability to reduce blue resazurin to red resorufin was measured at 540 nm. The average intensity of DMSO control was set as 100% alive and the percentage of intensity from each treated sample was calculated. The mean from biological triplicates was plotted with error bars representing the SEM in black. Nonlinear regression was analyzed and plotted in red, on which IC_50_ was based. (B) Growth curves of *B. subtilis* strain 168/WT in the presence of a range of HA concentrations. OD_600_ was measured every 30 min. HA was added at 2.5 h (arrow). The graph depicts the average and the SEM of biological triplicates. (C, D) No sporulation in response to HA. *B. subtilis* cells from a high-density overnight culture were treated with (C) HA (250 μg ml^-1^, 5 × MIC) or (D) DMSO (control), stained with FM4-64 and imaged by confocal fluorescence microscopy. Representative images are shown. Example spores in the DMSO control are indicated with arrows. Scale bar is 5 µm. (E) HA rapidly blocks respiratory chain activity in *B. subtilis*. Effect on respiratory chain activity measured by the reduction from blue resazurin to red resorufin at 540 nm. The average intensity of DMSO control in each group was set to 100% intensity. HA (250 μg ml^-1^, 5 × MIC) was added and after 5, 30 and 60 min, the percentage of conversion was calculated relative to the control. The mean from biological triplicates was plotted with error bars representing the SEM. The log 10 format was used for the scale of the Y axis. (F, G) HA treatment induced membrane permeability. (F) Membrane potential measurement. *B. subtilis* membrane potential levels were quantified using the fluorescent dye DiSC3(5). CCCP (2 μg ml^-1^, 5 × MIC), nisin (12.5 μg ml^-1^, 5 × MIC) or HA (250 μg ml^-1^, 5 × MIC) (as indicated) or DMSO was added after 5 min. The fluorescence was depicted as percentage of the value at the start (t = 0min) (y-axis) over time (x-axis, min). The mean from biological triplicates was plotted with error bars representing the SEM. (G) Membrane permeability measurement. Experiments were performed as in (F), except for using a different dye SYTOX-Green for quantifying membrane permeability.

To assess the toxicity of this compound on human cells, cytotoxicity assays were done using the HepG2 cell line, originating from human liver. The IC_50_ for HA was 226.6 μg ml^-1^ (Figure 1B), indicating that human cells have a higher tolerance for HA than Gram-positive bacterial cells.

To further investigate the effect of HA on cell lysis, we used the cell sporulation assay, which showed that following overnight HA treatment, only cell debris was observed (Figure 1C). In comparison, both intact cells and spores were shown in the control-treated cultures (Figure 1D). In addition, the result of the resazurin assay (Figure 1E) showed that after 5 min treatment, 99% of cells lost their viability. None of the cells seemed to be viable after HA treatment for 60 min. This suggested that HA has an immediate effect when added to bacteria. Taking these results into consideration, we hypothesized that HA might target the cell envelope.

### Pore formation is the initial effect of HA on bacterial cells

To investigate whether HA targets the cell envelope, we used DiSC3(5) to detect changes in transmembrane potential (20). Carbonyl cyanide *m*-chlorophenyl hydrazone (CCCP) and nisin were selected as positive controls, because they are known to destroy the membrane potential by disruption of cellular ionic homeostasis and generation of pores in the cell membrane, respectively (21, 22). Similar to treatment with CCCP or nisin, HA induced a rapid increase of fluorescence in cells (Figure 1F), suggesting immediate disruption of the membrane potential in response to HA.

To determine whether the loss of membrane potential was due to a pore-forming activity or not, cell permeability was investigated with SYTOX green (23). As shown in Figure 1G, the fluorescence intensity of the cell suspension gradually increased upon HA and nisin treatment, but not CCCP treatment. This suggests that the disruption of membrane potential in response to HA was caused by pore formation.

Vancomycin-and nisin-induced cell lysis is caused by binding to lipid II (22, 24). To determine if HA-induced pore formation was also mediated by binding to lipid II in the membrane, lipid I and lipid II were added to the bacterial cultures prior to addition of HA. If HA binds directly to these lipids, the externally added lipids will quench HA and hence, addition of HA will not affect bacterial growth (25). As positive controls, 1 × and 2 × MIC of vancomycin (0.17 and 0.35 μM) and nisin (1.86 and 3.73 μM) were used in this assay. *B. subtilis* cells were treated with antimicrobials in the presence or absence of different concentrations of these lipids and bacterial cell growth was determined (Table S3). As expected, the antimicrobial effects of nisin and vancomycin were quenched by exogenous lipid I and lipid II when the ratio of antimicrobials to lipids is less than 2:1. For HA, both 2 × MIC (258 μM) and 1 × MIC (129 μM) were tested in the presence of 10 μM or 150 μM of the different lipids. However, the activity of HA was not affected by addition of either of these lipids, suggesting the mechanism underlying pore formation by HA and nisin are distinct.

### Screening for HA-resistant mutants

Knowledge about antimicrobial resistance may provide insight into the underlying MoA. Inducing resistance by prolonged exposure to increasing concentrations of antimicrobial agents is a proven method. To obtain insight into the MoA of HA, we grew several colonies of *B. subtilis* strain 168 in the presence of 1 x MIC of HA in liquid medium for three days. Under these conditions, the liquid culture of three colonies grew and these were transferred sequentially to medium with 2-fold higher concentrations of HA. Over a period of 28 days, one culture was successfully grown in medium with 4 × MIC of HA. Resistance against an antimicrobial may be due to either gene mutation or gene expression adaptation. To distinguish between these possibilities, we cultured the resistant bacteria without HA for ten serial passages. Finally, HA resistance persisted in the M9015 strain, suggesting that resistance is caused by mutations in the genome.

The M9015 strain was not only resistant to HA, but the colonies also had a different appearance on agar plates than wild type. *B. subtilis* strain M9015 colonies had a more translucent appearance, but the colony size did not differ much between the wild type and M9015 strains (Figure 2A). We assumed that this interesting phenotype might be due to a slower growth rate (i.e. less cells), smaller cell size or a structural change in the cell wall (i.e. cells were more translucent). To investigate this, the growth curves of the strains were first compared. The growth pattern of both strains was similar, but the mutant had a 15 min longer period in the lag phase and a 0.2 lower final optical density at 600 nm in the stationary phase (Figure 2B). Next, the number of cells in the overnight culture was determined (Figure 2C, D). The cell density in the mutant strain was 2.2×10^8^ CFU ml^-1^, which was 57% higher than wild type (1.4×10^8^ CFU ml^-1^). Comparison of the cell size using a scanning electron microscope (Figure 2E, F) showed that M9015 mutant cells were 27 ± 5 % smaller than wild type cells (Figure 2G). These results suggest that the cell size affected the appearance of M9015 colonies.

**Figure 2.**
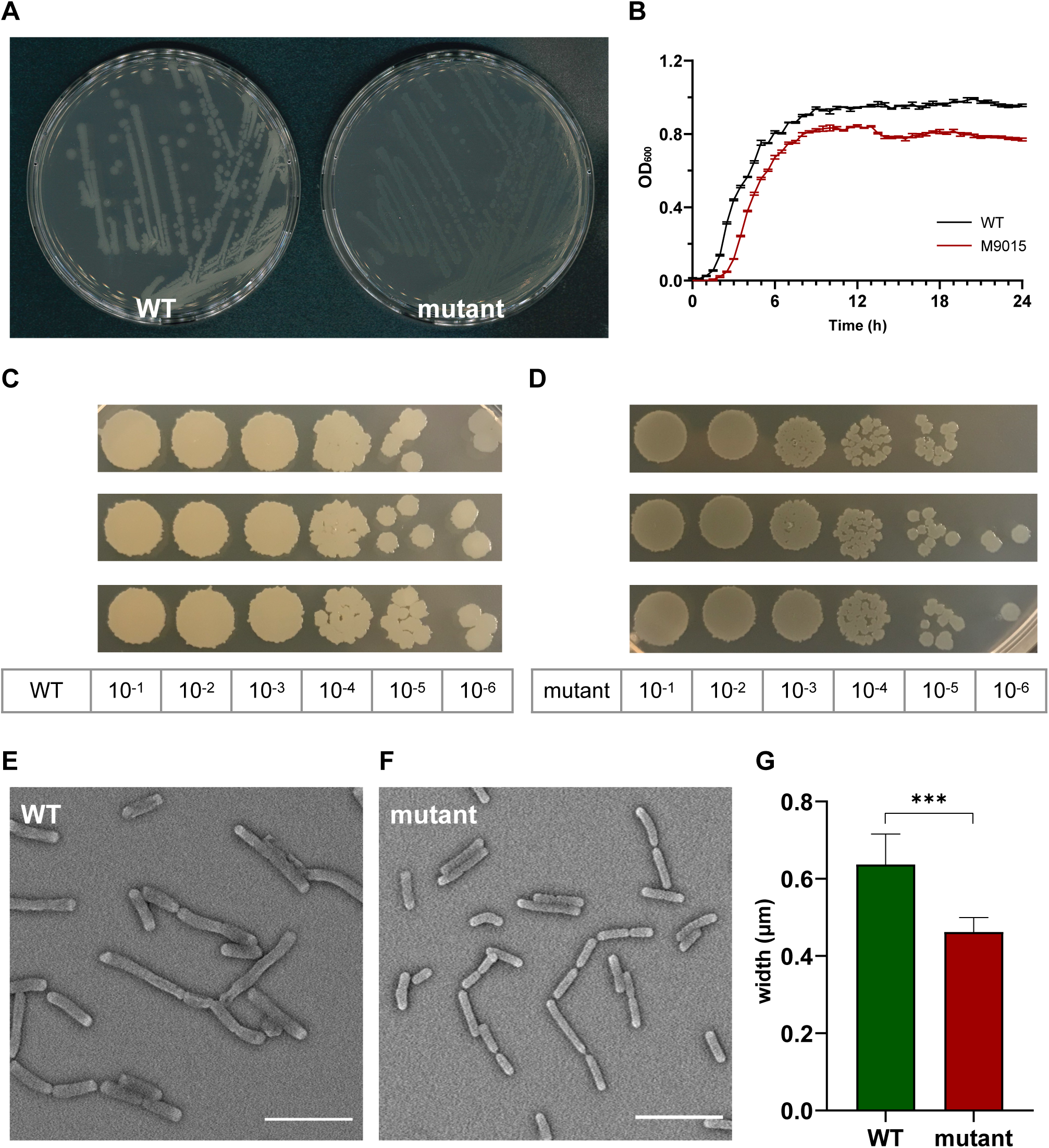
HA-resistant M9015 colonies are translucent and mutant cells are smaller. (A) *B. subtilis* strain 168/WT and M9015 plated on LB agar. (B) Growth curves of WT and M9015. OD_600_ was measured and the mean from biological triplicates was plotted with error bars representing the SEM. (C and D) CFU counting. The overnight cultures (18 h) of WT and M9015 were serially diluted, and 5 μL of each diluted sample was dropped on LB agar. Experiments were performed with biological triplicates. (E and F) The cells from overnight cultures (18 h) of WT and M9015 were fixed and imaged by scanning electron microscopy. Representative images are shown. The width of cells was measured and plotted in (G); mean with error bars representing the SEM is depicted. *** indicates p≤0.001 by student t-text.

To determine if there were changes in the structure of the cell wall that also contributed to the translucent phenotype, we analyzed the peptidoglycans. To this end, the cell wall was isolated from overnight cultures of both strains, followed by lysozyme digestion and HPLC analysis. Almost all peaks were matching between the M9015 mutant and wild type (Figure S2A). Peptidoglycan analysis of exponential phase cultures of these two strains showed similar results (Figure S2B). It is unlikely that the peptidoglycan composition of the cell wall contributed to the different appearance of the M9015 strain.

Cross resistance of the M9015 strain to other antimicrobials might provide valuable insight into the resistance class of this mutant strain. Therefore, we compared the MICs on M9015 mutant and wild type strains of antimicrobials from five major classes: nisin, vancomycin, chloramphenicol, moxifloxacin and rifampin. Interestingly, M9015 showed reduced sensitivity to rifampin, an RNA class antimicrobial, to an even higher extent than HA (4- to 8-fold; Figure 3). No significant differences were observed for any of the other antimicrobials, which included antimicrobials targeting the cell envelope. It is surprising that the HA-resistant M9015 strain is also more resistant to the RNA class antimicrobial rifampin, because our earlier experiments suggested that HA targets the cell envelope.

**Figure 3.**
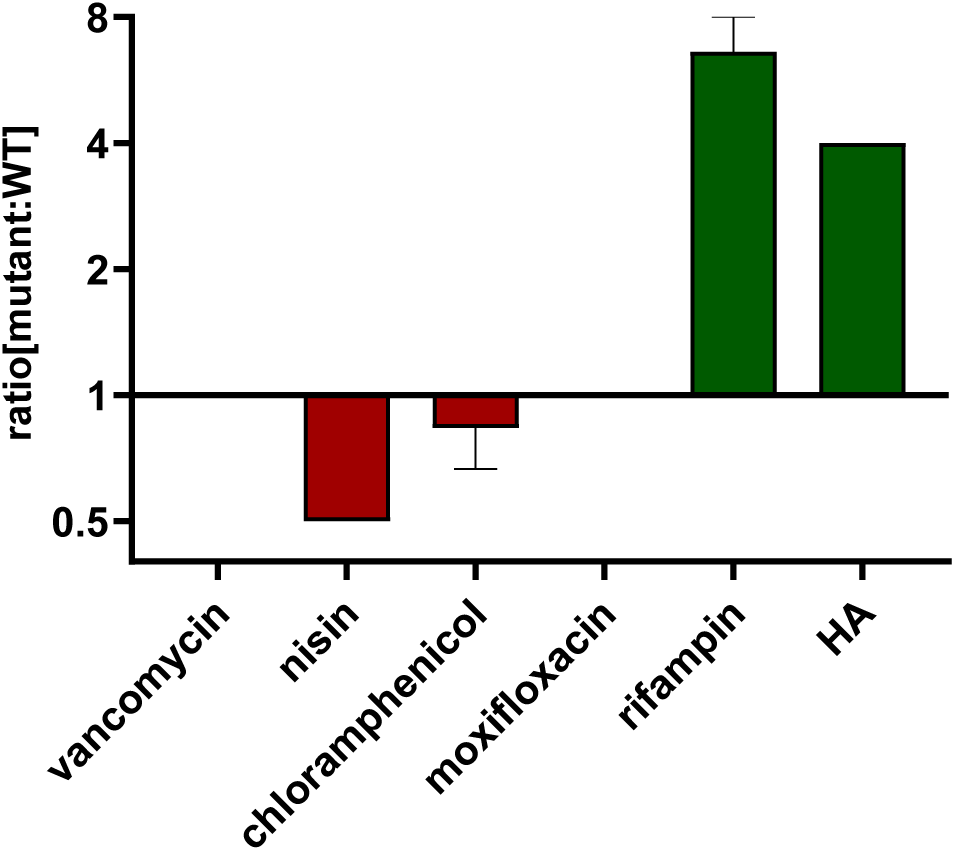
HA-resistant M9015 strain is also resistant to rifampin. MICs of selected antimicrobials against WT or M9015 were measured. The ratio of the MIC of a certain antimicrobial against M9015 to that against WT was calculated. The mean of the ratio from biological triplicates was plotted with error bars representing the SEM. The red bars indicated that M9015 was more sensitive to certain antimicrobial agents whereas the green bars indicated the opposite. The log 2 format was used for the scale of Y axis.

### Identification of mutations in M9015 by genomic sequencing

In order to determine which pathway(s) was/were affected in M9015, the genomes of M9015 mutant and wild type strains were sequenced by Next Generation Sequencing. The published genome sequence of *Bacillus subtilis* subsp. *subtilis* str. 168 was used as the reference genome. The mapping of the reads from our wild type strain against this referenced genome showed high coverage (only around 2,000 mutation sites), suggesting this reference is applicable, which facilitated the alignment of our sequencing data. Bioinformatic comparison of the wild type and mutant genomes resulted in the identification of ten potential variations (Table S4), five of which were suggested to be reliable by the Integrative Genomics Viewer (Figure S3, S4). These five mutations were located in the coding region of four different genes, *ymaB*, *flgL, atpE* and *yusO* (twice).

The mutations in *ymaB* and *atpE* were single-base substitutions causing an amino acid substitution in their protein products. YmaB is a putative Nudix hydrolase with RNA pyrophosphohydrolase activity (26), and the mutation we found resulted in a p.L138P substitution (Figure 4A). Although both these amino acids are non-polar and hydrophobic, their structures differ substantially, which might affect the activity of YmaB. The function of this gene is not clearly described in literature yet, and therefore the effect of the observed mutation remains unclear. The *atpE* gene encodes the subunit c of ATP synthase (27). In M9015, we identified a p.A51V substitution (Figure 4B). InterPro (https://www.ebi.ac.uk/interpro/) predicted the binding site of AtpE to be from p.33A to p.54E, indicating this mutation has a chance to be in the binding site. The property of these two amino acids is quite similar, but the larger size of Val in the mutant might affect the activity of AtpE severely.

**Figure 4.**
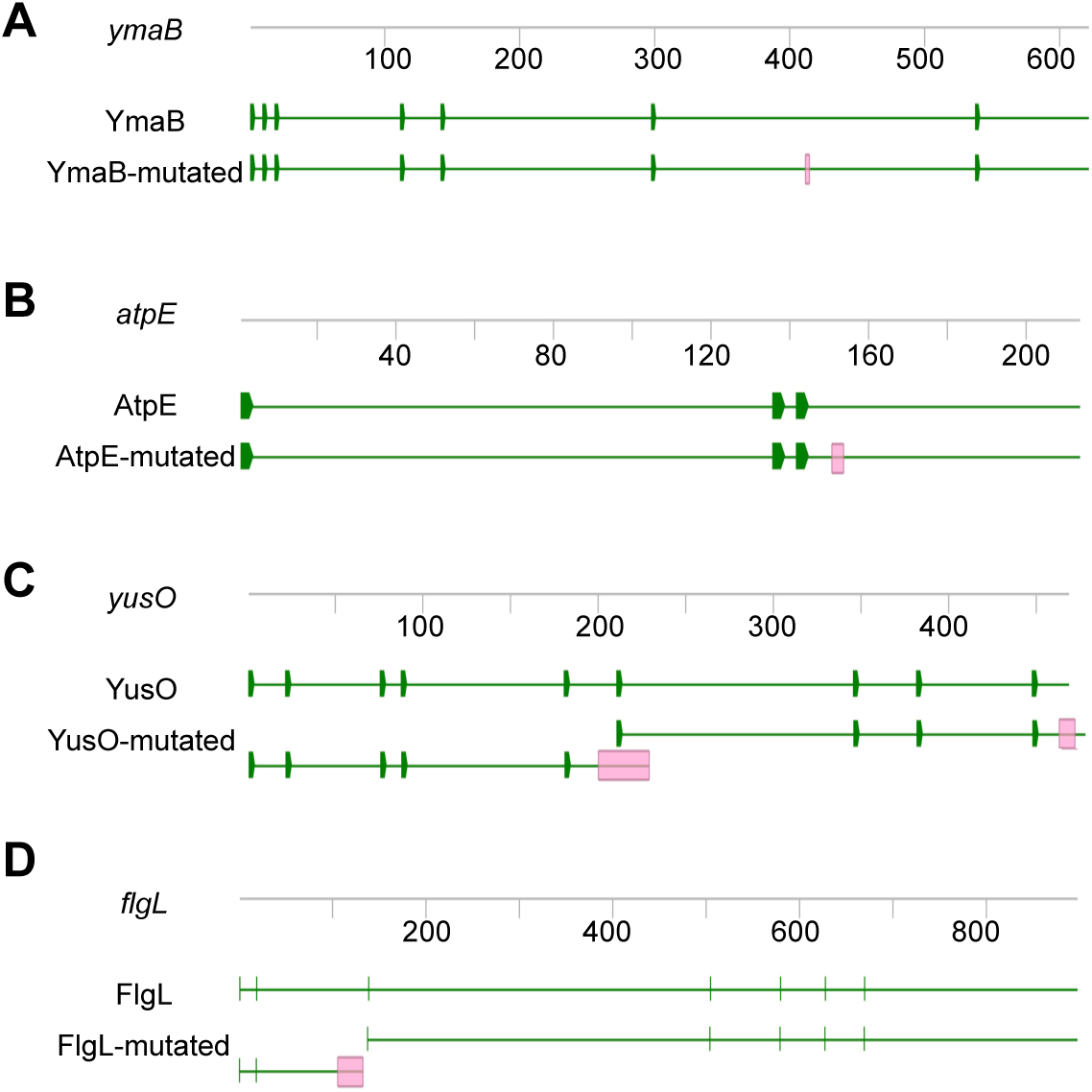
Mutated genes of M9015. The genes of *ymaB* (A), *atpE* (B), *yusO* (C) and *flgL* (D) are shown. Each gene was presented with DNA sequence (grey line with numbers indicating the length of DNA) and protein sequence (green line with green blocks indicating the sites of initiation codon). Red blocks in the mutated gene and protein sequences indicate the mutated sites.

The *yusO* gene expresses YusO, a transcription repressor, which binds to the yusOP promoter region and thus represses yusOP expression. YusP was reported to confer low-level resistance to fusidic acid, novobiocin, streptomycin, and actinomycin (28). This gene was disrupted by two insertions in M9015 mutant cells: a single-base insertion (c.199_200insA), introducing a frame shift and stop-codon and an in-frame nine-base insertion (c.454_455insGAGGAAACG). Therefore, the gene product would be different from the wild type strain (Figure 4C). Instead of expressing YusO, the M9015 mutant might produce two (non-functional) truncated proteins from this gene, dubbed YusO-m1 and YusO-m2. In YusO-m1, the frame shift would result in a premature stop p.A67D fsX10. There is an alternate initiation site just past the single base pair insertion in *yusO*, which might initiate the expression of YusO-m2. The nine base pair insertion near the 3’ end of *yusO* would lead to a three amino acids duplication near the C-terminus of YusO-m2 (p.G153_G155dup). However, whether YusO-m1 and/or YusO-m2 proteins are stably expressed in M9015 remains to be determined. Hence, it is not unlikely that mutation of *yusO* might contribute to HA resistance.

The *flgL* gene expresses FlgL, a flagellar hook-filament junction protein, which has a function in motility and chemotaxis (29). It had a one base pair deletion in M9015 cells, resulting in a truncated FlgL protein (p.K35SfsX9, FlgL-m1, Figure 4D). Potentially, a second FlgL protein, FlgL-m2 starting from Met47 of FlgL was produced in M9015 cells as well (Figure 4D). It has been suggested that disruption of this gene may lead to reduced motility, which might contribute to the translucent appearance of M9015 colonies as well.

To further study these gene functions, we ordered four mutants from Bacillus Genetic Stock Center (30), each contains a deletion of one of the four indicated genes (Table 1). The colonies of most mutants appeared normal, except for the Δ*atpE* strain, which had smaller and more translucent colonies (Figure S5). This indicates that the mutation in *atpE* is responsible for the translucent phenotype.

Next, we tested the MICs of HA and rifampin on these four mutants. As listed in Table 1, Δ*yusO* strain had a similar level of resistance to both HA and rifampin as our M9015 strain. In comparison, none of the other strains showed more resistance to either of the antimicrobials. This suggested that the disruption of *yusO* is the reason for HA resistance in the M9015 strain.

### HA may have multiple targets and only generate pores on the membrane at high concentration

Interestingly, since *yusO* is a transcriptional repressor of a multidrug efflux transporter (28), it is unlikely to confer resistance to an antimicrobial targeting the cell envelope. Nevertheless, we observed effects of HA on the cell envelope (Figure 1F, G). We generated two hypotheses regarding this finding. One is that HA might affect membrane integrity without direct targeting of the membrane. For instance, gentamicin exclusively targets ribosomes, which results in misfolded membrane proteins. Eventually, this will lead to membrane defects (31, 32). However, gentamycin did not induce changes in the membrane potential under conditions that membrane defects were induced, i.e treatment with gentamicin for up to 15 min (Figure S6). These results suggest that it is unlikely that HA affects pore formation by indirect mechanisms, such as production of misfolded proteins.

Our second hypothesis was that HA might have multiple targets, including a target in the membrane. This is reminiscent of triclosan, a multi-target antimicrobial that has specific targets in the membrane only at high concentrations (33). To investigate this hypothesis, we performed depolarization assays with different concentrations of HA. The results showed that HA did not affect cell membrane at 0.5 × MIC or 1 × MIC (Figure 5A). Only at 2 × MIC of HA, effects on the cell membrane were observed. Nisin treatment on the other hand showed effects on membrane polarization at all concentrations tested, even at 0.5 × MIC (Figure 5B). To investigate if *yusO* was required for HA-induced cell envelope damage, we tested cell permeability of mutant strains M9015 and Δ*yusO* with HA concentration of 100 μg ml^-1^, which is 1/2 × MIC for these two strains, but 2 × MIC on the wild type strain. As shown in Figure S7, pores were generated in both of the mutants, like in the WT control strain.

**Figure 5.**
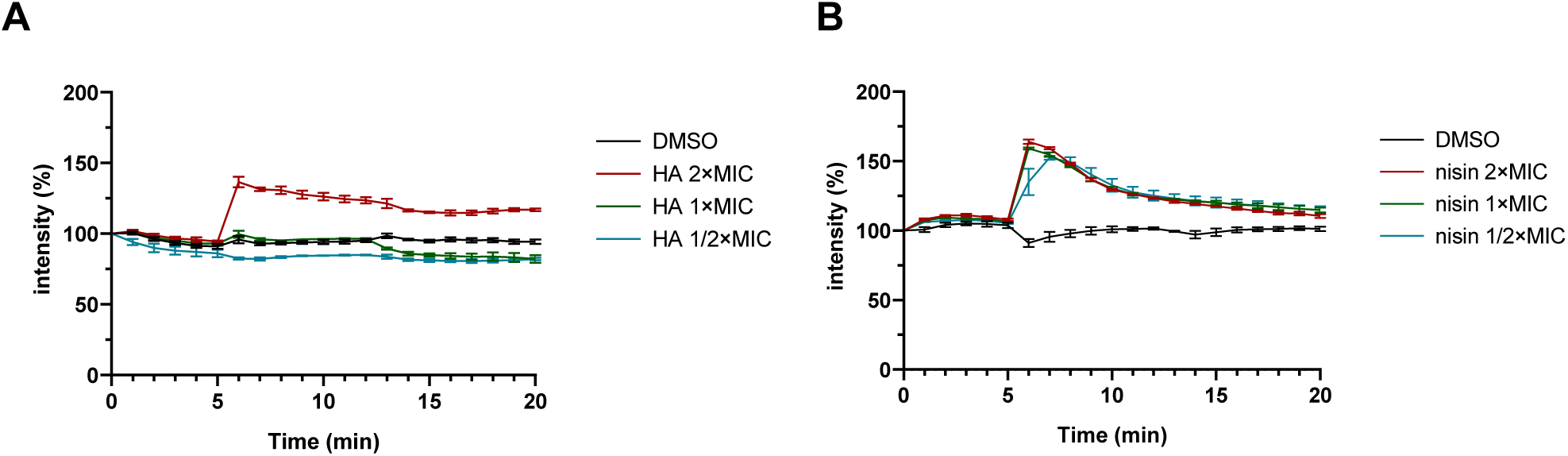
HA affects cell depolarization only at high concentration. *B. subtilis* membrane potential levels were quantified as in Figure 1F. Different concentrations of HA (A) or nisin (B) were added after 5 min. DMSO was used as control. The fluorescence was depicted as percentage of the value at the start (t = 0min) (y-axis) over time (x-axis, min). The mean from biological triplicates was plotted with error bars representing the SEM.

Altogether, these results suggest that HA is a multi-target antimicrobial with an intracellular target that remains to be identified and the cell membrane as target, but only at higher concentrations of HA.

## Discussion

Here, we describe the antimicrobial property of HA. The minimum concentration of HA to inhibit *B. subtilis* growth overnight was 50 μg ml^-1^, which was around 4 to 5 times less than the IC_50_ of human cells (Figure 1A, B). This suggested HA not to be very toxic. Treatment of bacteria with HA at concentrations below 50 μg ml^-1^ led to rapid arrest of bacterial growth and even lysis of the cells (a reduction in OD600). Interestingly, following treatment with 30 or 40 μg ml^-1^ HA, bacterial growth recovered 3 to 9 hours after the start of treatment, respectively (Figure 1B). The mechanism underlying recovery of the bacterial growth following treatment with HA below 50 μg ml^-1^ remains to be determined. The cells showed delayed but similar growth kinetics to control samples. HA may be degraded by the few live cells, or HA may be exhausted or quenched by the bacterial cells. The remaining live cells appear not to be affected by the previous presence of HA in the medium and are now able to grow freely again at the same growth rate as untreated cells.

Our results suggested that HA is a multi-target antimicrobial agent that generates pores in the cell membrane at high concentrations. Since HA did not bind to either of the provided lipids in our study (Table S3), the mechanism of the pore formation is still unclear. To further study the mechanism, other approaches, e.g. molecular dynamics simulations to model pore formation in lipid bilayers (34, 35), might provide insight into the underlying mechanism.

To further study the MoA of HA, we selected a low-level HA-resistant strain, *B. subtilis* strain M9015, by continuous exposure to HA. The M9015 strain contains four mutated genes, which has more translucent colonies than the wild type. We examined this phenotype in relation to HA-resistance. In the comparison of the growth patterns and cell size between mutant and wild type, we found that the size of M9015 cells was smaller. Smaller cell size might contribute to the translucent appearance of colonies, because the colonies will be thinner when the cells are smaller. The observed apparent reduction in growth might also be caused by reduced cell size, because the same number of bacterial cells will have a lower OD_600_ when the cells are smaller. The translucent phenotype Of the M9015 strain was similar in the Δ*atpE* strain, which lacks one of the four genes that are mutated in M9015. Given the similarity in phenotype, it is likely that the translucency is caused by lack of *atpE.* Loss of *atpE* is not involved in resistance to HA, because the Δ*atpE* strain showed similar sensitivity to HA as WT (Table 1). One of the four genes, *yusO*, confers similar resistance to bacteria as observed in the M9015 strain (Table 1). Interestingly, *yusO* has been implicated in antimicrobial resistance before (28). Yet, it has been reported that several point-mutations in *yusO* did not confer resistance to rifampin (28), which is not in line with our results. We found that both the HA-resistant strain M9015 and the Δ*yusO* strain had cross resistance to rifampin. In the previous work (28), they did not knock out the entire gene. In our M9015 strain and the Δ*yusO* strain, the function of *yusO* gene is totally disrupted, which might explain the discrepancy in observed resistance to rifampin.

Recently, HA was found to target acetohydroxyacid synthase (AHAS) in fungi (18). AHAS is the first enzyme in the branched-chain amino acid biosynthetic pathway. Since AHAS is also present in *B. subtilis* strain 168 (36), low concentrations of HA might target this enzyme in bacteria as well, thus affecting bacterial growth. To what extent inhibition of AHAS contributes to antimicrobial activity of HA towards bacteria remains to be determined.

To conclude, our results suggest that HA is a multi-target antimicrobial agent against Gram-positive bacteria. It targets the cell membrane, but only at high concentrations. We have developed a HA-resistant strain, M9015, and discovered that disruption of *yusO*, a component of the yusOP multidrug resistance operon was responsible for resistance to both HA and rifampin. The intracellular target of HA remains to be determined, but AHAS, the first enzyme in the branched-chain amino acid biosynthetic pathway is a good candidate.

## Materials and Methods

### Strains and reagents

*B. subtilis* strain 168 was used for MoA identification in this study (36). *O. flavum* (CBS 366.71) was obtained from the Westerdijk Fungal Biodiversity Institute (the Netherlands) and used for biologically active compound production. Pathogenic bacterial strains used for activity tests were either obtained from ATCC or they were clinical isolates (kind gift from University Medical Center Utrecht, the Netherlands) and they are listed in Table S2. *B. subtilis* mutants were obtained from Bacillus Genetic Stock Center (30) (Table 1). Commercial antimicrobials and resazurin were purchased from Sigma Aldrich. FM4-64, DiSC3(5) and SYTOX-Green were purchased from Thermo Fisher Scientific.

### Identification of HA

Identification of fungal SMs were performed as described before with minor modifications (14, 15). *O. flavum* was cultured on Malt Extract Agar (MEA) for 14 days. Secondary metabolites were extracted using ethyl acetate and separated using a Shimadzu preparative high performance liquid chromatography (HPLC) system with a C18 reversed phase Reprosil column (10 μm, 120 Å, 250 × 22 mm). The mobile phase was 0.1% trifluoroacetic acid in water (buffer A) and 0.1% trifluoroacetic acid in acetonitrile (buffer B). A linear gradient was applied of buffer B (5–95%) for 40 minutes. Fractions were collected and tested on *B. subtilis.* The active fraction was assessed for its purity through Shimadzu LC-2030 analytical HPLC using a Shimadzu Shim-pack GISTC18-HP reversed phase column (3 μm, 4.6 × 100 mm). LC-MS was performed on a Shimadzu LC-system connected to a Bruker Daltonics µTOF-Q mass spectrometer. High resolution mass spectrometry (HRMS) was measured on an LCT instrument (Micromass Ltd, Manchester UK). Elemental composition analyses were performed by Mikroanalytisch Labor Kolbe (Oberhausen, Germany). Finally, the compound was dissolved in 400 µL CDCl3 + 0,03% TMS and analyzed by Nuclear Magnetic Resonance (NMR) spectroscopy. More specifically, ^1^H-NMR, Hetronuclear Single Quantum Coherence (HSQC), Hetronuclear Multile-Bond Correlation (HMBC) and Correlation spectroscopy (COSY) spectra were measured at 600 MHz using a Bruker instrument. ^13^C-NMR was measured on the same instrument at 150 MHz.

### Microdilution assay

Minimum Inhibitory Concentration (MIC) was determined by broth microdilution assay as previously described (37). The freshly prepared early exponential-phase cell cultures of different strains were diluted 1 : 100 into Luria-Bertani (LB) medium, and then distributed in a 96-wells plate. Antimicrobials were tested starting at a 10 × dilution of the stock in DMSO, which was then serially diluted with a factor 2. MIC was defined as the lowest dilution at which bacteria did not grow, based on visual inspection after an overnight incubation at 37 °C.

### Growth curves

Overnight bacterial cultures were diluted 1 : 50 into fresh LB medium and incubated at 37 °C with shaking. OD_600_ of cultures were measured by a FLUOstar microplate reader (BMG Labtech) every 30 min for 24 h.

### Confocal microscopy

Microscopy was performed following the protocol described before with minor modification (15, 38). Briefly, samples were stained with indicated dyes (1.5 µM FM4-64 and/or 0.5 µM SYTOX-Green), immobilized on microscope slides covered with an agarose pad containing 1% agarose and LB medium, and imaged. Confocal microscopy was carried out using a Perkin Elmer UltraView VoX spinning disk microscope system. Experiments were done with biological triplicates. Images were analyzed using Fiji (39).

### Sporulation assay

This assay was applied as previously described with modifications (40). The freshly prepared early exponential-phase cell cultures of *B. subtilis* with an OD_600_ of 0.3 in LB media were centrifuged and resuspended in 1/10 of the original volume. Antimicrobials (5 × MIC) or DMSO (control) were added and 15 µL of the mixtures were transferred into 1.5 ml tubes to incubate with rolling at 37 °C for 5 h. Cells were then imaged as described above.

### Cytotoxicity assay

For the cytotoxicity assay, HepG2 cells were seeded in 96-well plates and grown in DMEM low glucose medium (ThermoFisher, 10567014) supplemented with 10% FBS. Test compounds were added in different concentrations, with a final concentration of 1% DMSO and cells were incubated for 20 h at 37 °C with 5% CO_2_. Next, resazurin was added to reach a final concentration of 0.1 mM. After 3 h incubation, the fluorescence was measured on a PHERAstar microplate reader (BMG Labtech) using an excitation wavelength of 540 nm and emission wavelength of 590 nm. The average intensity of DMSO control was set as 100% alive and the percentage of intensity from each treated sample was calculated. IC_50_ was calculated using nonlinear regression in GraphPad Prism. Experiments were conducted in biological triplicates.

### Resazurin assay

This assay was applied as previously described with modifications (38). Resazurin is an oxidation–reduction indicator (excitation at 530-570 nm, emission at 580-590 nm) (41). Freshly prepared early exponential-phase *B. subtilis* cultures with an OD_600_ of 0.3 in LB medium were treated with antimicrobials (5 × MIC) or DMSO (control) for 5, 30 and 60 min. Untreated cells and boiled cells (95°C for 10 min) were used to calculate the standard respiration and no respiration, respectively. Next, cells were washed with medium by centrifugation and afterwards resuspended in medium to adjust the OD_600_ to 0.15. Cells were then incubated with 30 µg ml^-1^ resazurin for 45 min at 37 °C. Absorbance of different samples was measured using a 540 nm/590 nm filter.

### Cell depolarization assay and cell permeability assay

DiSC3(5) is commonly applied for cell depolarization assay because it generally accumulates in well-energized cells. Disruption of membrane potential releases this probe from cells into the medium, resulting in an increase of overall fluorescence in the cell suspension (20). SYTOX green was used in the cell permeability assay. This dye is cell impermeable and its fluorescence signal increases significantly when bound to DNA (23). To perform these assays, the freshly prepared early exponential-phase cell cultures with an OD_600_ of 0.3 in LB medium were stained with DiSC3(5) or SYTOX green for 10 min. Then the fluorescence was measured on a PHERAstar microplate reader (BMG Labtech) using a 540 nm/590 nm filter for DiSC3(5), or a 485 nm/520 nm filter for SYTOX green. Experiments were conducted in biological triplicates.

### Lipid assay

This assay was applied as previously described with modifications (25). The appropriate amounts of antagonists lipid I, lipid II and C_55_-P in CHCl_3_ : MeOH (1 : 1) were added to the 96-well plates and the solvent was evaporated. The antagonists were re-dissolved in LB containing antimicrobials (2 × and/or 1 × MIC) or 1% DMSO in solution. The freshly prepared early exponential-phase cell cultures of *B. subtilis* were diluted 1 : 100 into LB medium, and then distributed in the 96-wells plate with antagonists and antimicrobials. After incubation overnight at 37 °C, bacterial growth was inspected visually.

### Screening for HA-resistant mutants

To obtain HA-resistant mutants, we grew *B. subtilis* in the presence of 1 x MIC of HA in LB medium at 37 °C with shaking. Once bacterial growth was observed, it was sequentially transferred to medium with 2-fold higher concentrations of HA. The whole process was set for a period of 28 days.

### Scanning electron microscopy

Overnight cultures of *B. subtilis* strain 168 and M9015 were transferred to fresh LB medium and grown to mid-log phase. Bacterial cells were then washed, fixed, dehydrated, mounted onto 12.5 mm specimen stubs and coated with gold to 1 nm as previously described (42). Samples were imaged using a Phenom PRO desktop SEM (Phenom-World BV). Cell width was measured from 10 cells for each strain from triplicate images using Fiji.

### Peptidoglycan analysis

This assay was performed following a previous protocol (43). The overnight bacterial culture of *B. subtilis* was diluted 1 : 50 into 400 ml fresh LB medium and incubated at 37 °C with shaking until reaching expected cell density. The cell culture was then chilled on ice and centrifuged at 5,000 x g for 10 min at 4 °C. After washing twice with cold PBS, the cell pellet was resuspended in 10 ml PB (25 mM sodium phosphate, pH 6.0), disrupted using beads beater, mixed with 10 ml of 10% (w/v) SDS in a boiling water bath, boiled for 30 min and incubated at 37 °C overnight. Next, the sample was centrifuged at 45,000 x g for 30 min at 20 °C to pellet the insoluble peptidoglycan, which was then washed with 10 ml of PB 4-6 times, suspended in 2 ml PB and transferred to a microcentrifuge tube. After adding 200 μg ml^-1^ of pronase and incubating for 2 h at 37 °C with agitation, TCA hydrolysis was performed at 4 °C for 18 h with constant shaking. Afterwards, the hydrolyzed sample was centrifuged at 45,000 x g for 30 min at 20 °C to pellet the insoluble peptidoglycan, which was then washed three times, suspended in 5 ml PB. After digestion of peptidoglycan using 0.5 mg ml^-1^ lysozyme overnight and reduction of sugars using sodium borohydride for 20 min, sample was ready for HPLC analysis, which was performed using a Shimadzu LC-2030 system with PDA detection (190-800 nm) and a Shimadzu Shim-pack GISTC18-HP reversed phase column (3 μm, 4.6 × 100 mm). The mobile phase was 50 mM sodium phosphate pH = 4.33 (buffer A) and 50 mM sodium phosphate pH = 5.1 with 15% (v/v) methanol (buffer B). A flow rate of 0.5 ml min^-1^ was applied using the following protocol: buffer A for 10 minutes followed by a linear gradient of buffer B (0– 100%) for 120 minutes, 100% buffer B for 10 minutes, another linear gradient of buffer B (100–0%) for 5 minutes and finally buffer A for 5 minutes.

### Genomic DNA sequencing

The bacterial cell wall of *B. subtilis* was lysed using 0.5 mg ml^-1^ lysozyme for 1 h at 37 °C. Next, genomic DNA was isolated using the Wizard Genomic DNA purification Kit (Promega) and sequenced by Illumina Next Generation Sequencing in University Medical Center Utrecht.

### Bioinformatic analysis

The raw sequencing reads were subjected to a panel of bioinformatic analyses including quality control using FastQC and Trimmomatic, alignment to reference genome using BWA-MEM algorithm and variant calling using Integrative Genomics Viewer (IGV). The published genome sequence of *Bacillus subtilis* subsp. *subtilis* str. 168 was used as the reference genome, which has a NCBI accession number of NC_000964.3.

## Acknowledgements

The authors would like to thank Anko de Graaff of the Hubrecht Imaging Centre for help with imaging, Ad Fluit of the Medical Microbiology department of UMC Utrecht for clinical bacterial isolates, Eefjan Breukink (Utrecht University) for Lipid II and derivatives, Ronnie Lubbers for critical reading of the manuscript and Marieke Kuijk for help with antimicrobial activity test. This project was supported by the Chinese Scholarship Council (CSC).

## Notes

### Competing Interest Statement

The authors have declared no competing interest.

