## Supplemental Material for "Harzianic Acid has multi-target antimicrobial activity against Gram-Positive Bacteria"

### Table S1: Assignments NMR-shifts, HMBC and COSY couplings for harzianic acid (in $CDCl_3$)

| | | Harzianic acid ($CDCl_3$) | | |
|---|---|---|---|---|
| # | δC[a] | δH[b] | HMBC[b] | COSY[b] |
| 1 | 13.7 | 0.95 (t), 3H | 2, 3 | 2 |
| 2 | 21.8 | 1.50 (m), 2H | 1, 3, 4 | 1, 3 |
| 3 | 35.5 | 2.24 (dd), 2H | 1, 2, 4, 5 | 3, 4 or 5[c] |
| 4 | 149.9 | 6.37[c] (m), 1H | 2, 3, 5, 6, 7[c] | 3, 6[c] |
| 5 | 129.6 | 6.38[c] (m), 1H | 2, 3, 6, 7[c] | 3, 6[c] |
| 6 | 147.6 | 7.55 (m), 1H | 4, 5, 7, 8 | 4 or 5[c], 7 |
| 7 | 119.1 | 7.00 (d, *J*=15.1 Hz), 1H | 5, 6, 8, 10 | 6 |
| 8 | 176.7 | - | - | - |
| 9 | 173.2 | - | - | - |
| 10 | 99.7 | - | - | - |
| 11 | 197.3 | - | - | - |
| 12 | 64.1 | 3.63 (dd, *J*= 10.6, 1,0 Hz), 2H | 9, 11, 13, 14, 18, | 13 |
| 13 | 33.8 | 1.89 (dd), 1H<br>2.48 (d), 1H | 11, 12, 14, 15, 19 | 12 |
| 14 | 79.9 | - | - | - |
| 15 | 36.0 | 2.02 (m), 1H | 13, 14, 16, 17, 19 | 16 or 17[c] |
| 16 | 17.5 | 0.99[c] (m), 3H | 14, 15[c] | 15 |
| 17 | 16.2 | 0.99[c] (m), 3H | 14, 15[c] | 15 |
| 18 | 26.6 | 2.97 (s), 3H | 9, 12 | - |
| 19 | 176.3 | - | - | - |

[a]= measured at 150 MHz, [b]= measured at 600 MHz, [c]= overlapping signals

**Table S2. HA MIC on pathogenic bacteria**

MICs of HA on different pathogenic bacteria were tested starting at a 400 μg $ml^{-1}$ which was then serially diluted with a factor 2.

| Strain | Gram | MIC (μg $ml^{-1}$) |
|---|---|---|
| *Acinetobacter baumannii* 1179 [a] | - | > 400 |
| *Acinetobacter nosocomialis* 14-8211 [a] | - | > 400 |
| *Enterobacter cloacae* complex MC04842 [a] | - | > 400 |
| *Escherichia coli* TEM-3 GVJS004 [a] | - | > 400 |
| *Klebsiella pneumoniae* SHV-18 GVJS006 [a] | - | > 400 |
| *Pseudomonas aeruginosa* ATCC57853 [b] | - | > 400 |
| *Stenotrophomonas maltophilia* GV20A226 [a] | - | > 400 |
| *Enterococcus faecium* VRE GV16D030 [a] | + | 100 |
| *Enterococcus faecium* GV15A623 [a] | + | 50 |
| *Listeria monocytogenes* GV21-4a [a] | + | 25 |
| *Staphylococcus aureus* MSSA 476 GVS0101 [a] | + | 50 |
| *Staphylococcus aureus* MRSA USA300 GVS1474 [a] | + | 200 |
| *Staphylococcus epidermidis* GV08A1071 [a] | + | 50 |
| *Streptococcus pneumoniae* 05A396 [a] | + | 25 |

[a] Gift from University Medical Center Utrecht;

[b] ATCC strains.

**Table S3. Exogenous Lipid II and its precursors did not affect HA antimicrobial activity**

Vancomycin, nisin and HA have an antimicrobial effect on *B. subtilis*. Quenching of this antimicrobial effect was tested by addition of exogenous Lipid I and Lipid II (either 1 μM or 10 μM for nisin and vancomycin; either 10 μM or 150 μM for HA) prior to the addition of indicated antimicrobials. Both 1 × and 2 × MIC of vancomycin (0.17 and 0.35 μM), nisin (1.86 and 3.73 μM) and HA (129 and 258 μM) were used. Bacterial growth was assessed and the assay scored as inhibited (-) or uninhibited (+) by the addition of exogenous lipid. NA means not tested.

| **Antagonist** | **Blank** | **Lipid I (1 μM)** | **Lipid II (1 μM)** | **Lipid I (10 μM)** | **Lipid II (10 μM)** | **Lipid I (150 μM)** | **Lipid II (150 μM)** |
|---|---|---|---|---|---|---|---|
| Vancomycin (2 × MIC) | - | + | + | + | + | NA | NA |
| Vancomycin (1 × MIC) | - | + | + | + | + | NA | NA |
| Nisin (2 × MIC) | - | - | - | + | + | NA | NA |
| Nisin (1 × MIC) | - | + | + | + | + | NA | NA |
| HA (2 × MIC) | - | NA | NA | - | - | - | - |
| HA (1 × MIC) | - | NA | NA | - | - | - | - |

**Table S4.** ***B. subtilis*** **strain M9015 harbors five mutations in four genes**

Bioinformatic analysis of the genome sequences of *B. subtilis* strain 168 and M9015 results in 10 possible mutations, five of which are reliable. The position of the mutations is indicated and is based on the reference genome NC_000964.3. The mutations were visualized using Integrated Genomics Viewer and the gene names of the verified reliable mutations are indicated.

| # Mutation | Position | WT | M9015 | Reliable? | Mutated gene |
|---|---|---|---|---|---|
| 1 | 353056 | AAGCAGCTGATC GAGCAGCTGA | AAGCAGCTGA | No | / |
| 2 | 1872540 | T | C | Yes | *ymaB* |
| 3 | 2152047 | A | C | No | / |
| 4 | 2480653 | CT | C | No | / |
| 5 | 2480666 | GT | G | No | / |
| 6 | 2581726 | GTTTTTT | GTTTTTTT | No | / |
| 7 | 3374690 | G | GA | Yes | *yusO* |
| 8 | 3374945 | AGAGGAAACGGA | AGAGGAAACGGAGGAAACGGA | Yes | *yusO* |
| 9 | 3638128 | ATTTTTTTT | ATTTTTTT | Yes | *flgL* |
| 10 | 3786681 | G | A | Yes | *atpE* |

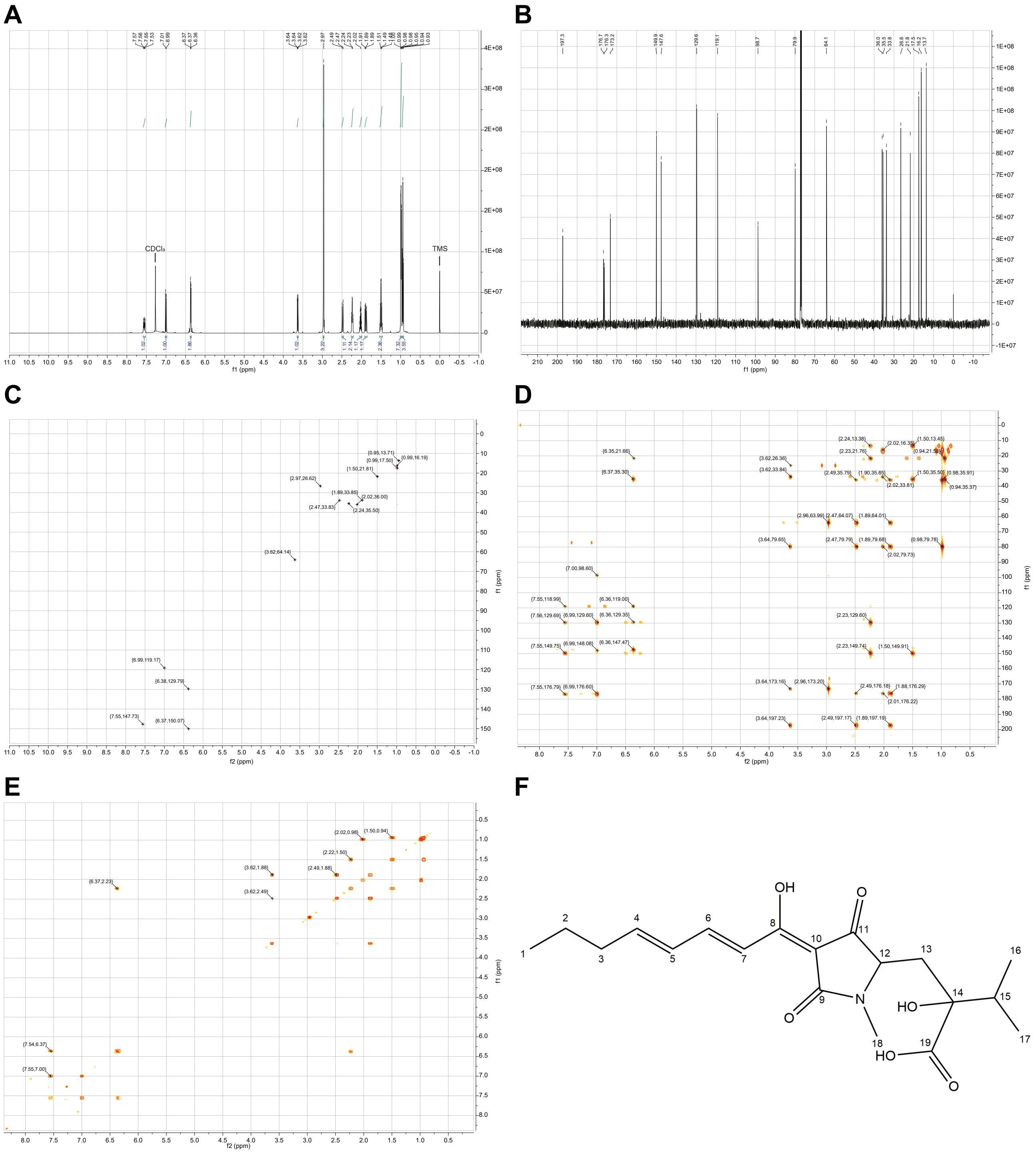


**Figure S1. NMR spectroscopy and chemical structure of harzianic acid.** (A) $^{1}$H-NMR spectrum, 600 MHz, $CDCl_3$. (B) $^{13}$C-NMR spectrum, 150 MHz, $CDCl_3$. (C) HSQC spectrum, 600 MHz, $CDCl_3$. (D) HMBC spectrum, 600 MHz, $CDCl_3$. (E) COSY spectrum, 600 MHz, $CDCl_3$. (F) Chemical structure of harzianic acid.

**A**

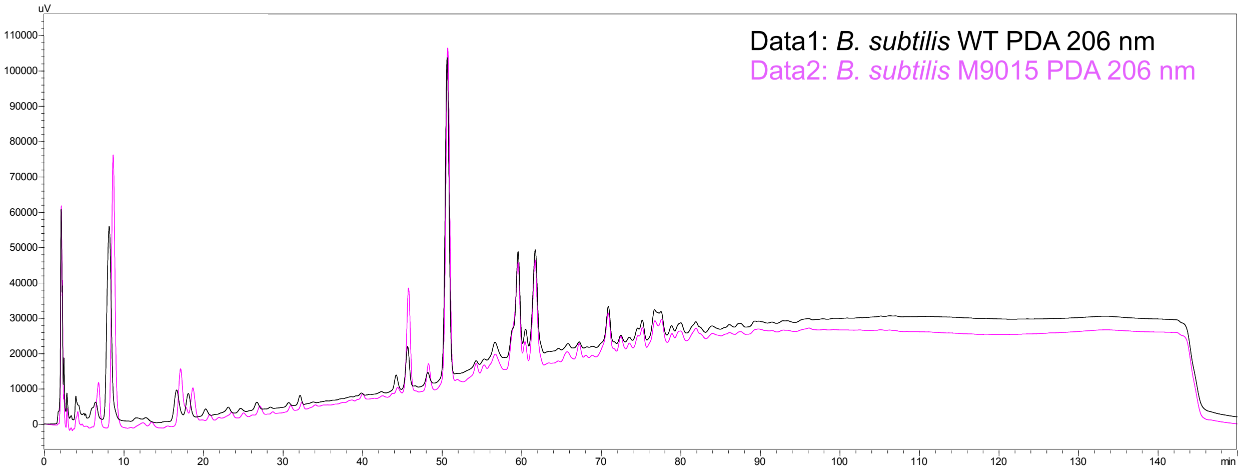


**B**

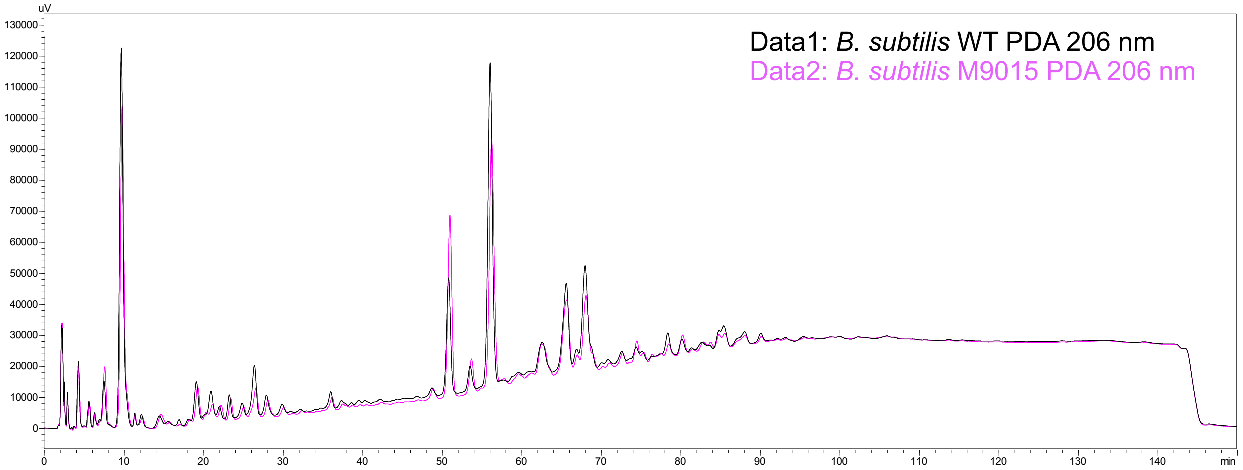


**Figure S2. Similar peptidoglycan pattern between *B. subtilis* strain 168 and M9015.** Peptidoglycan of both strains from overnight culture (A) and exponential phase culture (B) were isolated, digested and analyzed by analytical HPLC.

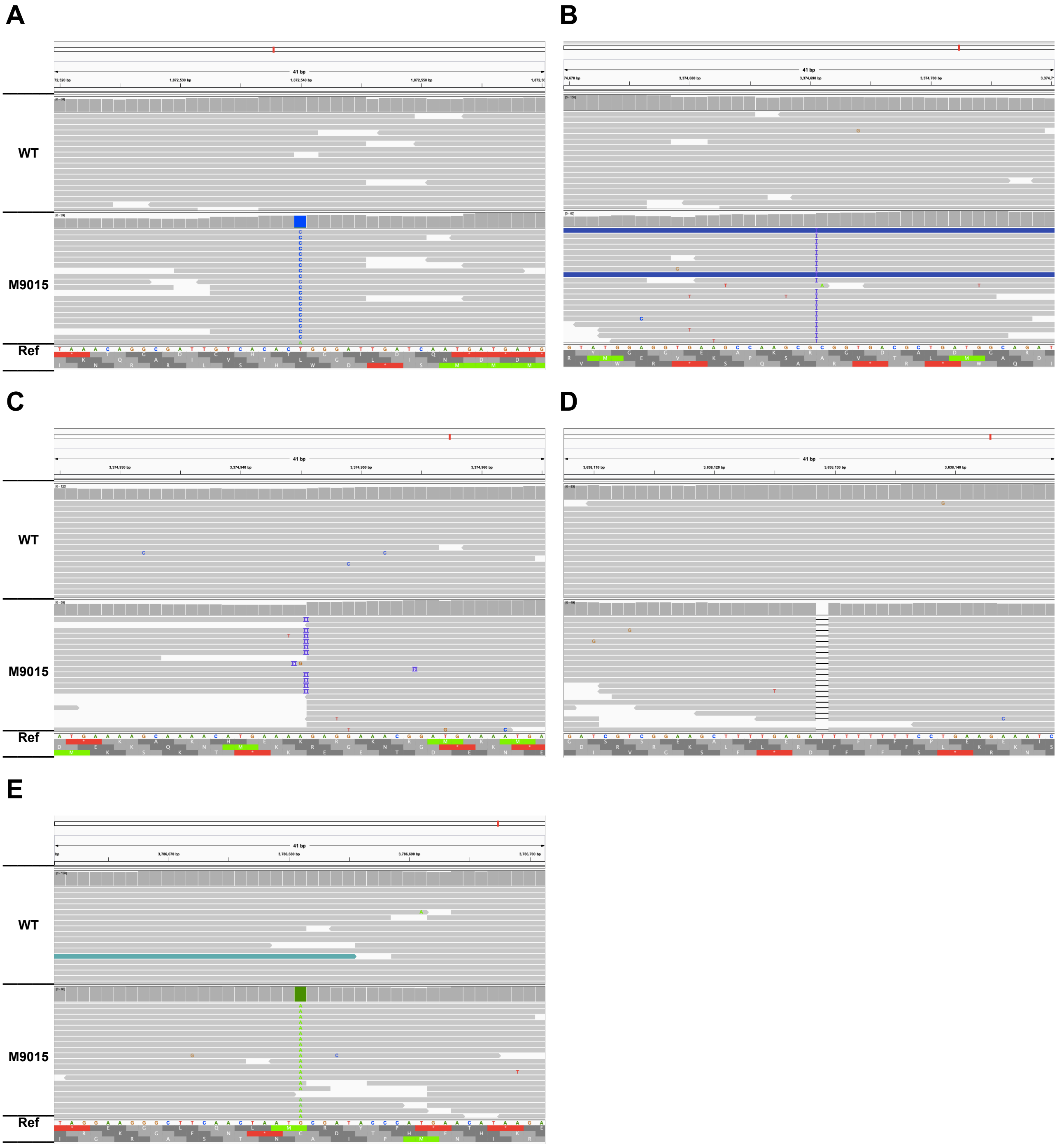


**Figure S3. Viewing gene variations with Integrated Genomics Viewer (I).** 10 predicted mutated sites were verified using Integrated Genomics Viewer. Five correctly predicted mutations (Mutation#2, #7-#10) (see Table 2) were listed in (A) to (E). “Ref” indicated the reference genome. Representative reads (around 15) were presented for both WT and M9015.

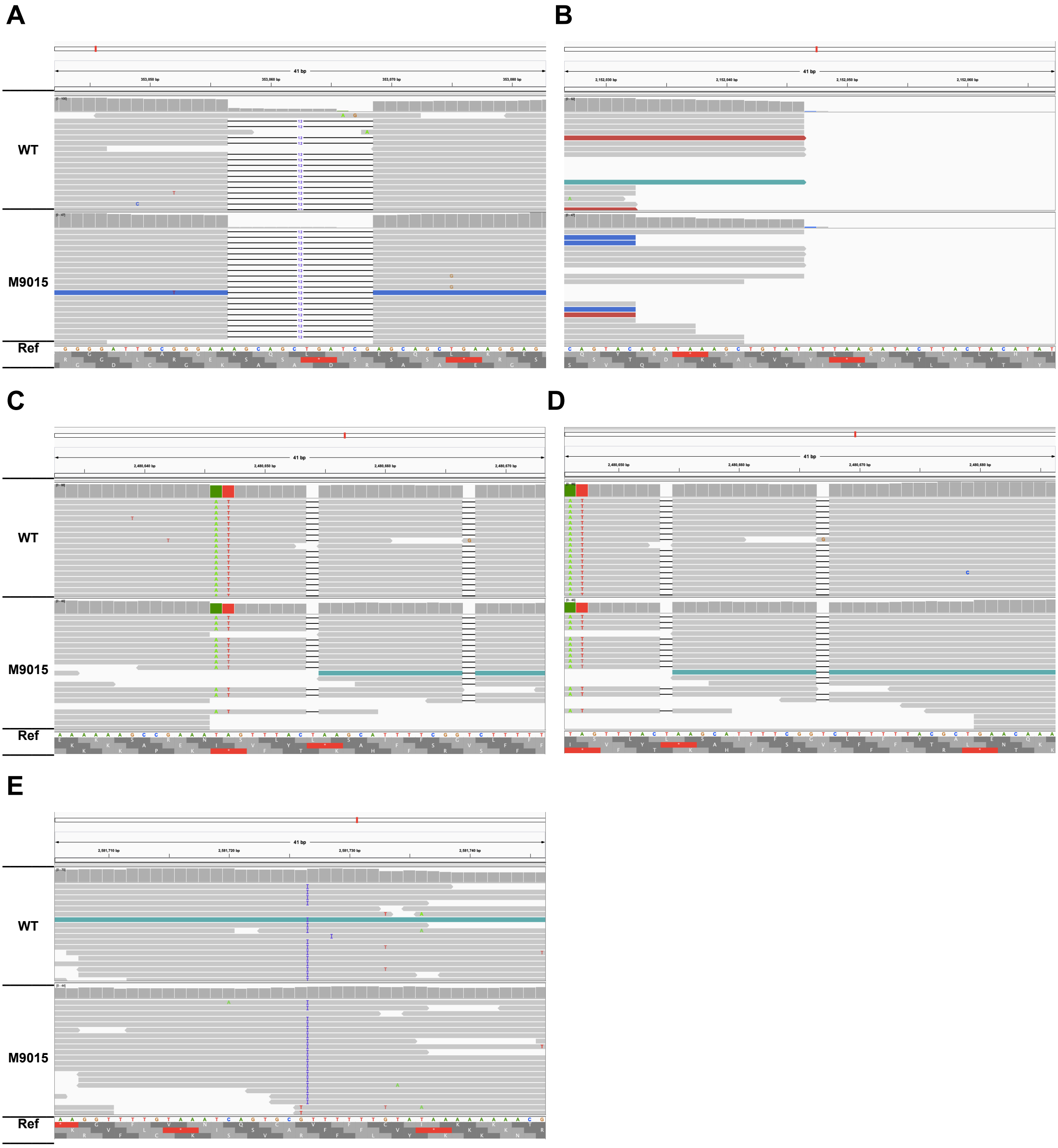


**Figure S4. Viewing gene variations with Integrated Genomics Viewer (II).** 10 predicted mutated sites were verified using Integrated Genomics Viewer. Five wrongly predicted mutations (Mutation#1, #3-#6) (see Table 2) were listed in (A) to (E). "Ref" indicated the reference genome. Representative reads (around 15) were presented for both WT and M9015.

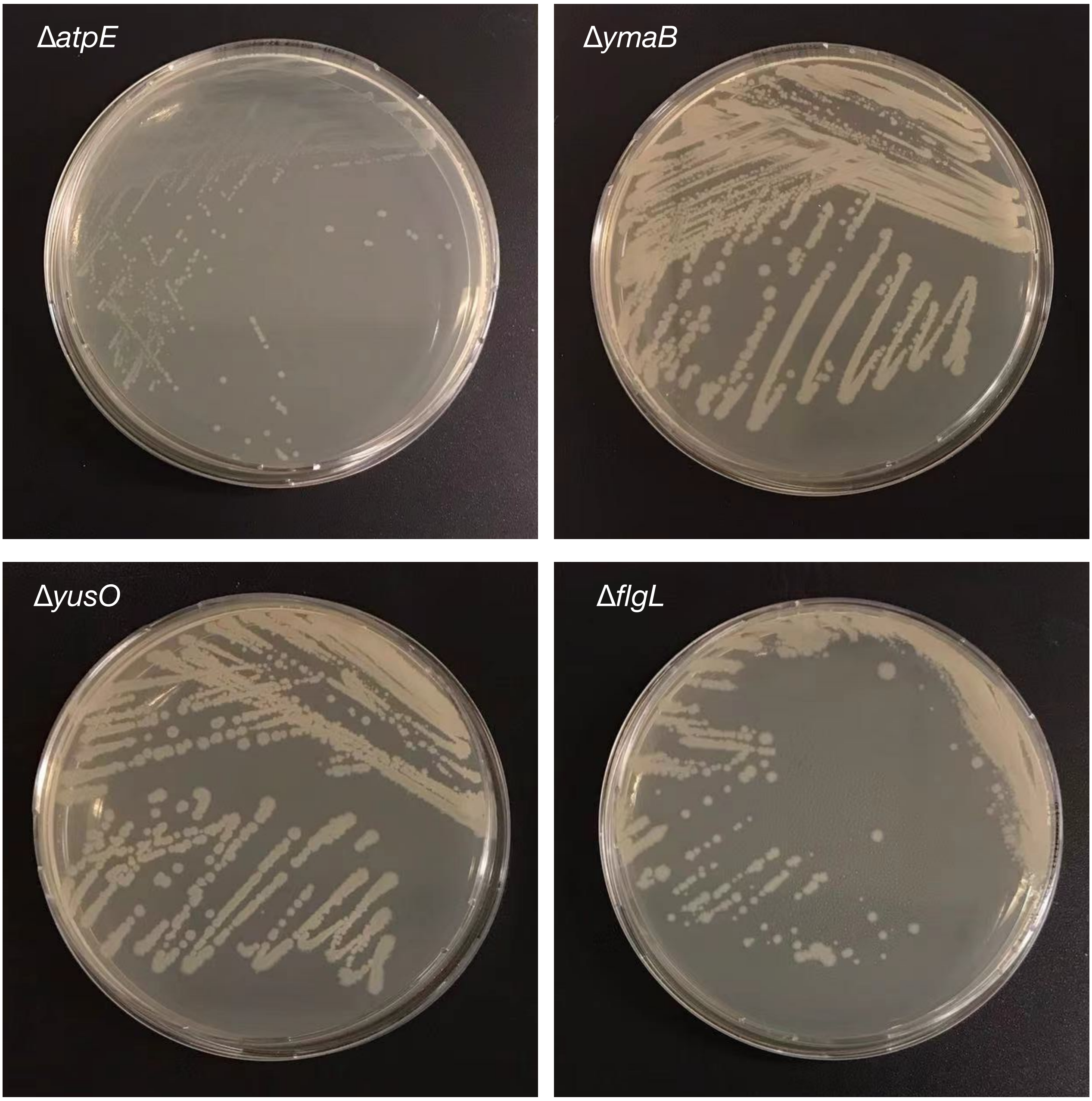


**Figure S5. Colony appearance of four mutants of *B. subtilis*.** *B. Subtilis* mutants were cultured on LB agar at 37 °C overnight. Plates were imaged.

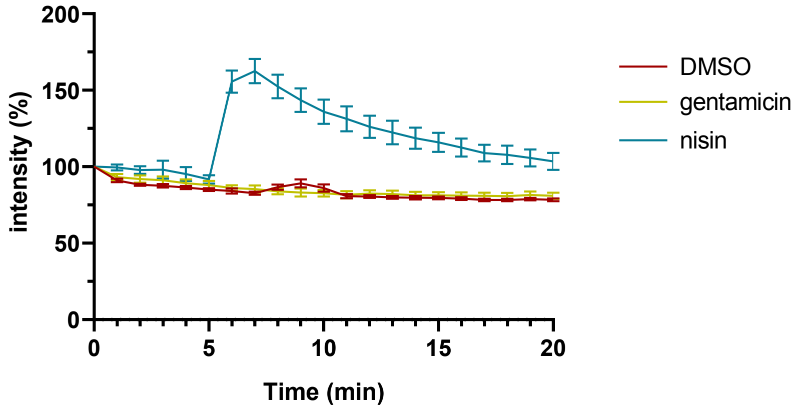


**Figure S6. Gentamicin did not affect cell depolarization.** *B. subtilis* membrane potential levels were quantified as in Figure 1F. Gentamicin, DMSO (blank control) or nisin (positive control) were added after 5 min. The fluorescence was depicted as percentage of the value at the start (t = 0min) (y-axis) over time (x-axis, min). The mean from biological triplicates was plotted with error bars representing the SEM.

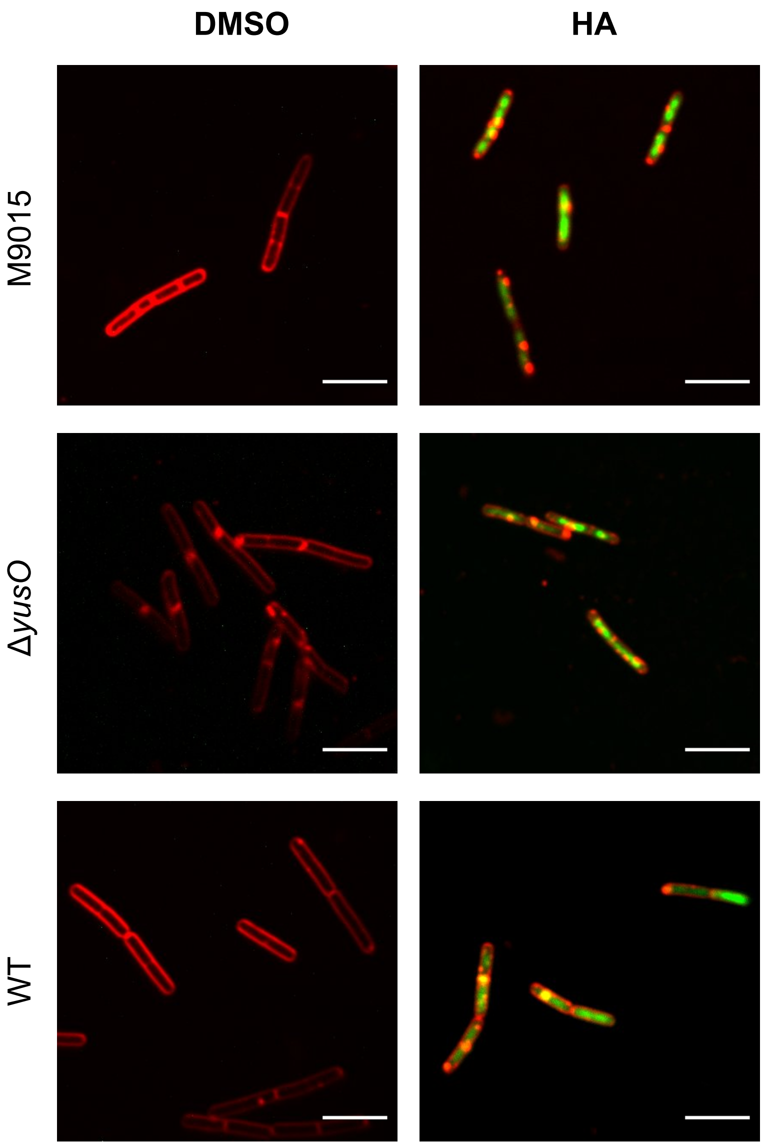


**Figure S7. Cell permeability determination of *B. Subtilis* strains.** *B. Subtilis* WT and mutants were stained with SYTOX-Green (green, nucleoid, cell-impermeable) and FM4-64 (red, cell membrane), treated with DMSO (control) or 100 μg $ml^{-1}$ HA for 15 min, and imaged. Representative cells are shown. Scale bar is 5 μm.
